# Discovery of five HIV nucleoside analog reverse-transcriptase inhibitors (NRTIs) as potent inhibitors against the RNA-dependent RNA polymerase (RdRp) of SARS-CoV and 2019-nCoV

**DOI:** 10.1101/2020.11.01.363788

**Authors:** Jialei Sun

## Abstract

The outbreak of SARS in 2002-2003 caused by SARS-CoV, and the pandemic of COVID-19 in 2020 caused by 2019-nCoV (SARS-CoV-2), have threatened human health globally and raised the urgency to develop effective antivirals against the viruses. In this study, we expressed and purified the RNA-dependent RNA polymerase (RdRp) nsp12 of SARS-CoV and developed a primer extension assay for the evaluation of nsp12 activity. We found that nsp12 could efficiently extend single-stranded RNA, while having low activity towards double-stranded RNA. Nsp12 required a catalytic metal (Mg^2+^ or Mn^2+^) for polymerase activity and the activity was also K^+^-dependent, while Na^+^ promoted pyrophosphorylation, the reverse process of polymerization. To identify antivirals against nsp12, a competitive assay was developed containing 4 natural rNTPs and a nucleotide analog, and the inhibitory effects of 24 FDA-approved nucleotide analogs were evaluated in their corresponding active triphosphate forms. Ten of the analogs, including 2 HIV NRTIs, could inhibit the RNA extension of nsp12 by more than 40%. The 10 hits were verified which showed dose-dependent inhibition. In addition, the 24 nucleotide analogs were screened on SARS-CoV primase nsp8 which revealed stavudine and remdesivir were specific inhibitors to nsp12. Furthermore, the 2 HIV NRTIs were evaluated on 2019-nCoV nsp12 which showed inhibition as well. Then we expanded the evaluation to all 8 FDA-approved HIV NRTIs and discovered 5 of them, tenofovir, stavudine, abacavir, zidovudine and zalcitabine, could inhibit the RNA extension by nsp12 of SARS-CoV and 2019-nCoV. In conclusion, 5 FDA-approved HIV NRTIs inhibited the RNA extension by nsp12 and were promising candidates for the treatment of SARS and COVID-19.

## Introduction

Severe acute respiratory syndrome coronavirus (SARS-CoV) and 2019-nCoV (SARS-CoV-2) belong to *Betacoronavirus* genus, *Coronaviridae* family and they have a single-stranded, positive-sense RNA genome which is approximately 30kb [1, 2]. SARS-CoV originates from bat and is transmitted to human through an intermediate host, palm civet [3, 4]. 2019-nCoV is also widely accepted to have a bat origin as it is highly similar to a bat coronavirus RaTG13 throughout the genome with 96.2% identity [5]. The high identity suggests a close relationship between the two viruses and there is no recombination event for both of them after they separated from each other evolutionally [4]. However, it is still unclear whether and when an intermediate host have been involved in the zoonotic transmission to human. Soon after the outbreak, pangolin was proposed to be the intermediate host when a pangolin-CoV was discovered which shared 91.02% identity with 2019-nCoV on whole genome level [6, 7]. The discovery provides evidence for the presence of an intermediate host, but 91.02% identity is still too low to suggest a direct transfer from the pangolin to human. Recently, it is reported that ACE2 from multiple animals can bind the spike protein of 2019-nCoV and support infection [8, 9], especially the ACE2 from monkeys which is identical to and binds as efficient as human [8, 10]. These studies also provides evidence for the involvement of an intermediate host. In addition, studies have revealed that 2019-nCoV has been under different types of natural selection. Its genome contains uncommon high ratio of neutral mutations which are featured by the predominance of C-U substitution [11-13]. The neutral mutations suggested there was strong selection pressure on 2019-nCoV and raised the possibility of that the virus stayed in a single type of host, possibly bat, for a very long period. The spike of 2019-nCoV appears to have been optimized for the binding to human ACE2 with a 1000-fold higher binding affinity than its closest relative, RaTG13 [14, 15]. And the spike also has acquired a polybasic furin cleavage site which is essential for its entry into host cell [16, 17]. These studies reveal strong purifying selection on the spike region of 2019-nCoV [18, 19], suggesting the adaptation from its original host to human is a rapid process. To explain the natural selection on 2019-nCoV, several hypotheses have been proposed, including selection in an animal host before zoonotic transfer, selection in humans following zoonotic transfer and selection during passage [14]. To better understand the origin of the virus, more studies are required.

For antiviral development, currently, there are at least three promising viral drug targets: spike protein (S), main protease (Mpro)/3C-like protease (3CLpro), and RNA-dependent RNA polymerase (RdRp). The spike of SARS-CoVs is a trimer and has two subunits, S1 and S2. The binding to its human receptor angiotensin-converting enzyme 2 (ACE2) is mediated by the receptor binding domain (RBD) located in the S1 subunit [20-23]. Upon binding to ACE2, spike undergoes conformational change to facilitate the fusion between viral membrane and host cell membrane [24]. The binding affinity between 2019-nCoV spike and ACE2 was reported to be 10-fold and 1000-fold higher than the spikes of SARS-CoV and its closest known relative RaTG13, respectively [15, 21]. High receptor binding affinity must have enhanced the human-to-human transmission of 2019-nCoV. As an essential factor for the infectivity of SARS-CoVs and the main target of neutralizing antibodies upon infection, spike has been a major drug target for antiviral discoveries, including neutralizing antibodies development such as LY-CoV555 and JS016 [25-30], inhibitor development [31] and vaccines [32-36]. According to Covid-19vaccinetracker.org, by September 2020, 35 vaccines in total were under different stages of clinical trials for COVID-19 and 10 of them were 2019-nCoV spike or spike subunit-based [37].

The main protease of SARS-CoVs, Mpro, also called 3CLpro, is encoded by nsp5 region. It is the major enzyme for polyprotein cleavage to produce mature and functional viral proteins [38]. Due to its essential role for SARS-CoVs, Mpro has been an attractive target for antiviral drug discoveries [39]. It is a cysteine protease and needs to form homodimer to be active [40-43]. The substrate residues at the cleavage site are conservative [44]. Currently, there are at least three drug discovery strategies for Mpro: structure-based drug design, biochemical assay screening and virtual screening. Since the outbreak of COVID-19, intense efforts have been made on virtual screening and multiple drugs have been predicted to be effective [45-50]. Drugs targeting Mpro are usually classified into two subgroups: peptidomimetics and small molecules [51]. Lopinavir/ritonavir (LPV/r), repurposed from HIV protease inhibitors, belongs to the peptidomimetics group. Although LPV/r was predicted to be potent initially, no benefit was reported after it was systematically investigated for the treatment of COVID-19 in clinical trials [52-57],

SARS-CoVs have two RNA-dependent RNA polymerases (RdRp): nsp12 and nsp8. Nsp12 is the main RdRp and plays the major role in the replication of RNA genome [58, 59]. Nsp8 is a non-canonical, sequence-specific RdRp and proposed to be the primase for SARS-CoVs [60, 61]. To replicate long RNA, nsp12 needs to interact with nsp7 and nsp8 to form a stable replication complex [62, 63]. Like all viral RdRps, nsp12 is right hand-shaped and composed of a fingers domain, a palm domain and a thumb domain [62]. The RNA polymerase activity is mediated by seven conserved motifs (A to G) [64, 65]. In addition, distinguished from most viruses, the nsp12 of SARS-CoVs also contains a nidovirus-unique N-terminal extension domain (NiRAN) which can covalently bind guanosine and uridine and has nucleotidylation activity [66]. The function of NiRAN in RNA replication is still unclear. As an attractive target for antiviral discoveries, efforts have been made on nsp12 since the outbreak of COVID-19, which mainly focus on drug-repurposing of FDA-approved drugs, especially on nucleotide analogs. Approaches for the repurposing include virtual screening, *in vitro* biochemical assays and cell culture-based assays [67, 68]. Virtual screening usually focuses on the binding between nsp12 and a compound, although interfering the interface between nsp12 and nsp8 or nsp7 has also been investigated [69, 70]. Hundreds of hits have been yielded by virtual screening, such as remdesivir, ribavirin, galidesivir, tenofovir, IDX-184 [71], and sofosbuvir [71-73]. Biochemical assays take advantage of purified proteins, mostly nsp12/nsp7/nsp8 replication complex, and have also identified useful antiviral nucleotide analogs against 2019-nCoV, including sofosbuvir, alovudine, zidovudine, tenofovir, abacavir, lamivudine, emtricitabine, cidofovir, valganciclovir/ganciclovir, stavudine and entecavir [72, 74-77]. These nucleotide analogs could be incorporated into extending RNA and terminate further chain elongation. Cell culture-based assays involve infectious live viruses and are the most direct approach to identify antivirals among the three. Studies using this approach have identified ribavirin, remdesivir, gemcitabine, and tenofovir in disoproxil fumarate form (TDF) to be inhibitors for SARS-CoVs [78-82]. Among them, ribavirin and remdesivir have been evaluated in clinical trials [83, 84] and remdesivir has been approved by FDA recently as it can shorten the time to recovery for patients.

In this study, we would focus on drug-repurposing of FDA-approved nucleotide analogs using a novel biochemical assay. And we managed to use nsp12 itself, instead of nsp12/nsp7/nsp8 complex, as the sole target for antiviral discovery. The nsp12 of SARS-CoVs was expressed and purified and its RNA extension activity was characterized under various conditions, especially its dependence on K^+^ and Mg^2+^. Furthermore, we developed a competitive assay which was able to quantify the relative inhibition abilities among nucleotide analogs, instead of using previously reported chain termination assays which could only give qualitative identification whether a nucleotide analog was a chain terminator for nsp12. Using the competitive assay, we identified 10 nucleotide analogs with >40% inhibition on SARS-CoV nsp12. Two of the hits were HIV NRTIs. We further evaluated all 8 FDA-approved HIV NRTIs and discovered 5 of them could inhibit the nsp12 of SARS-CoV and 2019-nCoV.

## Materials and Methods

### Chemicals

6-Mercaptopurine-TP (NU-1110S), 6-chloropurine-TP (NU-1109S), clofarabine-TP (NU-874), 6-methylthio-GTP (NU-1130S), 6-thio-GTP (NU-1106S), stavudine-TP (NU-1604S), 8-oxo-GTP (NU-1116S), gemcitabine-TP (NU-1607S), acyclovir-TP (NU-877), ganciclovir-TP (NU-275S), lamivudine-TP (NU-1606L) and zidovudine-TP (NU-989S) were purchased from Jena Bioscience. Tenofovir-DP (FT44596), 2’-C-M-GTP (NM08170), 2’-C-M-CTP (NM29280), emtricitabine-TP (ME16706) and gemcitabine-TP (ND09708) were purchased from Carbosynth. 2’-O-methyl-UTP (N-1018), 2’-azido-2’-dUTP (N-1029), 2’-amino-2’-dUTP (N-1027), ara-UTP (N-1034), 3’-O-methyl-UTP (N-1059), 2’-F-2’-dUTP (N-1010-1) and didanosine-TP (N-4017-1) were purchased from Trilink. Remdesivir-DP was synthesized by WuXi AppTech. Sofosbuvir-DP (HY-15745) was purchased from MedChemExpress. Ribavirin-TP (sc-358826) was purchased from Santa Cruz Biotechnology. Zalcitabine-TP (Z140050) and abacavir-TP/carbovir-TP (C177755) were purchased from TRC Canada.

### RNA primer-template annealing

RNA primer and templates were synthesized by Genscript. The sequences were modified from a previous study [85] and designed to be non-secondary structure forming. To form an RNA primer-template (P/T) complex, the primer (Cy5.5-5′-AACGUCUGUUCGCAAAAAGC-3′) was mixed with a template (5’-CUUAUUCGAGCUUUUUGCGAACAGACGUU-3’) in a ratio of 1:3 in 50 mM NaCl. The mixture was heated up to 95°C and slowly cooled down to 25°C by a PCR machine. The program was: 95°C for 5 min, 2°C decrease/10 s for 23 cycles, 51°C for 5 min, 2°C decrease/20 s for 14 cycles. The RNA primer was annealed to a stem-loop forming template (5’-CUAUUGACUUGCUUUUUCGCUACAGACGUU-3’) by the same procedure.

### Template-dependent RNA primer extension assay

Nsp12 template-dependent primer extension activity was determined by annealed RNA primer-template (P/T) complex. The assay condition contained 25 mM Tris HCl pH 8.0, 50 mM KCl or NaCl, 1 mM DTT, various concentrations of MgCl_2_ or MnCl_2_, 10 nM P/T complex, 32 nM purified nsp12 and 100 μM rNTPs (25 μM each). 10% Glycerol, 5 mM NaCl and 0.02% triton X-100 were introduced by nsp12 stock. Extra 1 mM NaCl was introduced by P/T complex which was prepared in 50 mM NaCl. The reaction was performed in 25 μl system and incubated at 37°C for 2 h. Poly A assay was performed in a similar condition containing 25 mM Tris HCl pH 8.0, 50 mM KCl or NaCl, 1 mM DTT, 0.5 mM MgCl_2_, 10 nM P/T complex, 16 nM nsp12 and 250 μM either ATP, UTP, GTP or CTP or a combination. Water was used as control. Combination assay of nsp12, nsp7 and nsp8 was also performed in a similar condition: 25 mM Tris HCl pH 8.0, 1 mM DTT, 0.5 mM MgCl_2_, 10 nM P/T complex, 100 μM rNTPs and 50 mM KCl. Concentrations of nsp12, nsp7 and nsp8 were 16 nM, 400 nM and 400 nM, respectively. Reaction was quenched by adding 50 µl quenching buffer which contained 90 mM Tris base, 29 mM taurine, 10 mM EDTA, 8 M urea, 0.02% SDS and 0.1% bromophenol blue. The reaction was then denatured at 95°C for 20 min, analyzed by 15% denaturing polyacrylamide urea gel electrophoresis and visualized using an Odyssey scanner (LI-COR).

### Combinational assay of nsp12 and helicase nsp13

The assay condition contained 25 mM Tris HCl pH 8.0, 50 mM KCl, 1 mM DTT, 1 mM MgCl_2_, 10 nM P/T complex, 16 nM nsp12, 210 or 420 nM nsp13, and 100 μM rNTPs (25 μM each). 10% Glycerol, 5 mM NaCl and 0.02% triton X-100 were introduced by nsp12 and nsp13 stocks. Extra 1 mM NaCl was introduced by P/T complex stock. The reaction was performed in 25 μl system and incubated at 37°C for 2 h. Activity on single-stranded RNA in the assay was performed with 10 nM the RNA primer instead of P/T complex. The concentrations of nsp12, nsp13, nsp7 and nsp8 were 16 nM, 420 nM, 6.25 μM and 1.25 μM, respectively.

### Back-priming extension assay

The activity of nsp12 in this assay was determined by the synthesized Cy5.5-labeled single-stranded RNA primer without a template. The assay condition contained 25 mM Tris HCl pH 8.0, 50 mM KCl, 1 mM DTT, 1 mM MgCl_2_, 10 nM RNA primer, 16 nM purified nsp12 and 100 μM rNTPs (25 μM each). 5% or 10% glycerol, 2.5 or 5 mM NaCl, and 0.01% or 0.02% triton X-100, were introduced by nsp12 stock, depending on the volume of nsp12 stock added in the system. Extra 1 mM NaCl was introduced by RNA primer stock which was prepared in 50 mM NaCl. The assay was performed in 25 μl system reaction at 37°C for 30 min, unless otherwise specified.

### Competitive assay for nsp12 and nsp8 screening

The inhibitory effects of 24 FDA-approved nucleotide analogs against nsp12 and nsp8 were determined using the exact back-priming extension assay mentioned above which was performed in 25 μl system and contained 16 nM purified nsp12 and 100 μM rNTPs (25 μM each). Nucleotide analogs were evaluated at 4 mM except for emtricitabine-TP which was evaluated at 2 mM due to low stock concentration. 4 mM ATP, UTP, GTP or CTP were used as controls. Assay with 100 μM rNTPs only was also included as control. The screening assay for nsp12 was incubated at 37°C for 30 min. Nsp8 was screened at 1.25 μM and incubation period for nsp8 was extended to 60 min due to the relatively lower activity of nsp8. Verification of nsp12 hits and hits evaluation on 2019-nCoV nsp12 were performed in the same condition. Reaction system was 25 μl and 15 μl, respectively. Extension products were then analyzed by 15% denaturing Urea-PAGE gel electrophoresis and scanned by an Odyssey scanner (LI-COR). Extended products were quantified using Image Studio software (LI-COR) and the intensity of ATP, UTP, GTP and CTP products were averaged. Product intensity of nucleotide analogs were then compared to the rNTP average of which the activity was defined as 100%, relative intensity and percent of inhibition were calculated.

### Concentration optimization of catalytic metals for nsp12 activity

The assay condition contained 25 mM Tris HCl pH 8.0, 50 mM KCl, 1 mM DTT, 10 nM RNA primer, 16 nM nsp12 and 100 μM rNTPs. The assay also contained 5% glycerol and 0.01% triton X-100 which were introduced by nsp12 stock. 3.5 mM NaCl was introduced by nsp12 stock and RNA primer stock. For catalytic metals optimization, nsp12 activities were determined at various concentrations of MgCl_2_ or MnCl_2_, from 0.0625 to 2 mM. Control was performed with same volume of water. The assay was re-performed with adding 0.5 mM EDTA into the reaction system, to chelate and block the catalytic activity of endogenous metals. For EDTA titration of endogenous metals, nsp12 activity was determined at various EDTA concentrations (1 nM to 100 μM) without adding MgCl_2_ or MnCl_2_. Reaction mix was incubated at 37°C for 30 min.

### Concentration optimization of KCl and NaCl for nsp12 activity

The assay condition contained 25 mM Tris HCl pH 8.0, 1 mM MgCl_2_, 1 mM DTT, 10 nM Cy5.5-labeled RNA primer, 16 nM nsp12 and 100 μM rNTPs. The assay also contained 10% glycerol, 0.02% triton X-100 and 6 mM NaCl introduced by nsp12 and Cy5.5-RNA stocks. Nsp12 activity was determined at various concentrations of KCl or NaCl, from 10 to 200 mM, with water used as control. An enzymatic control was performed with 50% glycerol instead of purified nsp12. Reaction mix was incubated at 37°C for 30 min.

### Time-course study of nsp12

The activity of nsp12 was determined in the same condition as back-priming extension assay. The reaction mix was prepared on ice and then incubated at 37°C. At various time-points (5-120 min), reactions were quenched and denatured immediately. The reaction products were then visualized and analyzed. Reaction with 50% glycerol instead purified nsp12 was used as enzymatic control which was incubated at 37°C for 120 min.

### Protein purification

All sequences in this study were synthesized by Genscript and constructed into pMAL-c5x vector with N-MBP and C-6X His tags. The sequences of SARS-CoV nsp12, nsp7 and nsp8 were adopted from a previous study [62] and constructed into pMAL-c5x vector between Nde I and EcoR I restriction sites. SARS-CoV nsp13 (nt 16167 to 17966, accession no.: NC_004718.3) was constructed into the vector between Nde I and BamH I sites. And 2019-nCoV nsp12 (nt 13468 to 16233, accession no.: NC_045512.2) was constructed between Nde I and Pst I. For protein purification, p-MAL-c5x-nsp12-His was transformed into *E. coli* BL21 (DE3) (Transgen, CD601-02) and protein expression was induced by an auto-induction system (71757-5, EMD Millipore). Upon overnight induction at 25°C, bacterial cell culture was collected and lysed with lysis buffer (20 mM Tris pH 8.0, 500 mM NaCl, 10% glycerol, 0.5% triton X-100, 2 mM MgCl_2_, 0.1 mM ZnCl_2_, 10 mM DTT, and 0.5X protease inhibitor cocktail, HY-K0010, MCE). Cell lysis was centrifuged at 5000 g for 30 min at 4°C to remove cell debris. Supernant was then applied to amylose resin (E8021L, NEB) for 3-5 h at 4°C to bind MBP-nsp12. Resin was then washed three times with washing buffer (20 mM Tris pH 8.0, 500 mM NaCl, 10% glycerol, 1% triton X-100, 1 mM DTT). To release nsp12 from MBP tag, the resin was treated with factor Xa protease (P8010L, NEB) overnight at 4°C. Supernant was collected and applied to a Ni-NTA resin (88222, Thermo) for 1-2h at 4°C to bind nsp12. The resin was washed three times with washing buffer and nsp12 was eluted with elution buffer (20 mM Tris pH 8.0, 50 mM NaCl, 10% glycerol, 0.1% triton X-100, 1 mM DTT, 300 mM imidazole). The concentration of glycerol was then adjusted to 50% and nsp12 stock was stored at −20°C. Nsp7, nsp8, nsp13 and 2019-nCoV nsp12 was purified by the same procedure. The concentrations of purified proteins were quantified by running an SDS-PAGE and comparing them with serially diluted BSA.

## Results

### Protein purification of the nsp12 of SARS-CoV and 2019-nCoV

Proteins in this study were expressed in *Escherichia coli* with MBP at N-terminus and 6X His at C-terminus. MBP was removed by factor Xa protease to produce His-tagged proteins for downstream assays. To simplify, his tag was omitted from the protein names. **Figure 1A** showed the purified SARS-CoV and 2019-nCoV proteins on SDS-PAGE and the targets to purify were indicated by stars **(*)**. The identities of proteins were confirmed by Mass spectrometry **(Supplementary Table 1)**. Production of SARS-CoV nsp12 per liter was low compared to nsp7, nsp8 and nsp13. Nsp12 had a large size (100 KD) which must have affected its correct folding during expression. The bulky size must also have restricted the access of factor Xa protease to its cleavage site which was located between MBP and nsp12, leading to the decrease of cleavage efficiency. These could explain the low production observed for nsp12. The production of 2019-nCoV nsp12 was low as well, which was consistent with SARS-CoV. In addition, as shown in the figure, there were 5 extra protein bands for nsp12 which were **(a)** MBP-nsp12, **(b)** 60 KD chaperone protein GroEL, **(c)** MBP, **(d)** elongation factor Tu, and **(e)** 30S ribosomal protein S3, as identified by Mass. GroEL is a major chaperone for protein folding in *E. coli* [86], it might have bound to nsp12 during expression to promote and maintain the correct folding of nsp12. Production of nsp7, nsp8 and nsp13 was relatively abundant and nsp8 showed an alternative product at 10 KD, indicated by **(#)**, which should be due to non-specific cleavage by factor Xa protease. The alternative product was confirmed by Mass as well.

**Figure 1.**
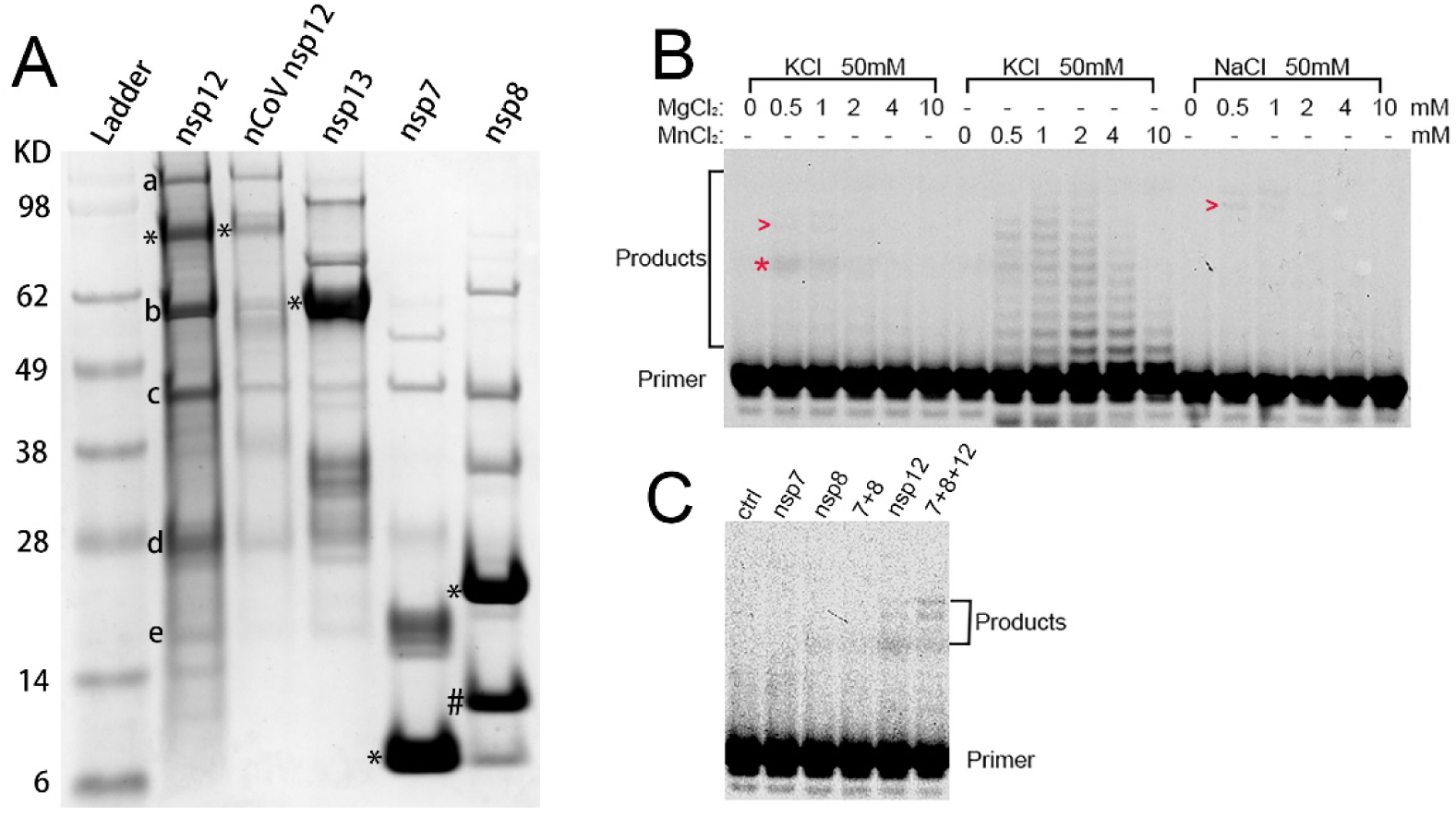
Protein purification and template-dependent RNA primer extension activity of SARS-CoV nsp12. **(A)** Purified nsp12, nsp13, nsp7 and nsp8 of SARS-CoV and nsp12 of 2019-nCoV were determined by SDS-PAGE and indicated by stars **(*)**. Nsp8 alternative cleavage by factor Xa protease was indicated by **(#)**. Five impurities of SARS-CoV nsp12 **(a-e)** were determined by Mass spectrometry. **(B)** The activity of SARS-CoV nsp12 on a double-stranded primer-template RNA complex was evaluated by primer extension assay at various concentrations of MgCl_2_ or MnCl_2_ combined with 50 mM either NaCl or KCl. The assay contained 25 mM Tris HCl pH 8.0, 1 mM DTT, 100 μM rNTPs, 32 nM nsp12 and 10 nM Cy5.5-labeled RNA primer which was annealed to its template. Reaction mix was incubated at 37°C for 2h and analyzed by denaturing Urea-PAGE. The gel was scanned and extended products were indicated by red **(*)** or **(>). (C)** The activity of nsp12 were also evaluated in combination with nsp7 and nsp8. Concentrations of nsp12, nsp7 and nsp8 were 16 nM, 400 nM and 400 nM, respectively. 50% Glycerol was used as control.

### SARS-CoV nsp12 had low extension activity on double-stranded RNA

It has been reported that the RNA polymerase activity of SARS-CoV nsp12 is primer dependent [59]. Therefore, upon purification, a Cy5.5-labeled RNA primer was annealed to its RNA template to form an RNA primer-template (P/T) complex and the activity of nsp12 was determined using the P/T complex. The condition for activity determination contained 50 mM KCl or NaCl combined with various concentrations of MgCl_2_ or MnCl_2_. Active nsp12 would extend the primer and fully extended product would have 9 more nucleotides (nt). As shown by **Figure 1B**, consistent with a previous study, nsp12 only showed activity with distributive product bands in the presence of Mn^2+^, while very low activity was observed for Mg^2+^ [58]. However, as Mg^2+^ had a higher concentration and was the predominant catalytic metal for DNA/RNA polymerases intracellularly [87], we did not continue the study with Mn^2+^ which did not represent the real physiological condition. For Mg^2+^, the activities of nsp12 were evaluated for both K^+^ and Na^+^. Both showed very weak activities with faint bands indicated by red **(>)**. As extension lengths of the bands were close to the predicted 9 nt, we initially thought they were real template-dependent primer extension activities. However, when we further performed studies with ATP only instead of all 4 rNTPs, nsp12 showed same extension products **(Figure S1)**. This activity was not detected for UTP, GTP, CTP or their combinations. Therefore, nsp12 could utilize ATP as the sole source for primer extension under the conditions tested and the activities observed for Mg^2+^ was poly A activity, not template-dependent primer extension. To identify the real activity, we then determined the activity of nsp12 in the presence of nsp7 and nsp8 **(Figure 1C)**, as nsp12 has been reported to form complex with nsp7 and nsp8 to be active. As compared to nsp12 alone, the combination with nsp7 and nsp8 had similar level of products, suggesting that nsp7 and nsp8 did not significantly enhance the activity of nsp12 in the condition tested. Nsp7, nsp8 and combination of the two were not active completely. Taken together, these results suggested nsp12 had low activities towards the double-stranded RNA.

### Nsp12 had high activity towards single-stranded RNA

Despite of repeatings under various conditions for months, we were not able to improve the activity with the double-stranded RNA P/T complex, possibly due to the low concentration of nsp12 (16 nM) which was much lower than reported studies [59, 88]. Then we noticed that, in the presence of K^+^, there was another weak product band **(Figure 1B)**, which was indicated by red star **(*)**. The product had extended 7 nt as compared to the distributive bands catalyzed by Mn^2+^. As the product was weak and 2 nt shorter than fully extended product which was 9 nt, we hypothesized that the activity of nsp12 was hindered by the helix force of double-stranded RNA and it needed to couple with helicase to achieve high activity and long extension. To test, we purified the helicase nsp13 of SARS-CoV **(Figure 1A)** which had been reported to have protein-protein interaction with nsp12 [89-91]. Following the purification, we designed a stem loop-forming P/T complex as well as the P/T complex used above with perfect base-pairings for the determination of nsp12 activity in combination with nsp13 **(Figure 2A)**. The molar ratios of nsp12 and nsp13 in the combination were 1:13 and 1:26. Reaction with 50% glycerol was used as control. As shown by **Figure 2B**, upon combination, the activity of nsp12 was enhanced by nsp13 in a dose-dependent manner. The enhancement was especially obvious for the P/T complexes with stem-loop structure. This result suggested nsp13 did enhance the activity of nsp12. However, extension length of the product was still 7 nt instead of the predicted 9 nt. This observation was against the coupling model between nsp12 and nsp13, as the helix unwinding by nsp13 should lead to longer extension of nsp12. Then, we realized that nsp12 was active towards the single-stranded RNA primer. The enhancement of nsp12 activity observed was due to the release of more single-stranded primer by nsp13 from the double-stranded P/T complex. To confirm, the assay was performed with the single-stranded primer without the annealing to a template and almost full activity was observed for nsp12 **(Figure 2B)**. This type of activity of nsp12 should be due to the back-priming of RNA primer of which the 3’ end bends back and pairs within the primer to form an extendable hairpin structure, which has been reported for nsp12 previously [63, 88]. Taken together, these results showed that nsp12 had high activity towards the single-stranded RNA primer. In addition, nsp8 and nsp7+nsp8 complex also showed high activities which was not surprising as nsp8 was presented to be the primase for SARS-CoVs [60, 92]. Interestingly, nsp13 and nsp7 also showed weak activities which possibly had relationship with their abilities to bind RNA. As shown in **Figure 1B**, the product had 7 nt extension, we analyzed the primer sequence (Cy5.5-5′-AACGUCUGUUCGCAAAAAGC-3′) and found that the 5’-AGC-3’ at 3’-terminus could form a double parings with 3’-UUG-5’ in the sequence. The full extension for this paring was 7 nt (5’-AGACGUU-3’) which had 2 A, 2 G, 2 U and 1 C **(Figure 2C)**. Therefore, the sequence of the extended product was predicted to be Cy5.5-5′-AACGUCUGUUCGCAAAAAGCAGACGUU-3′, which formed a stem-loop structure.

**Figure 2.**
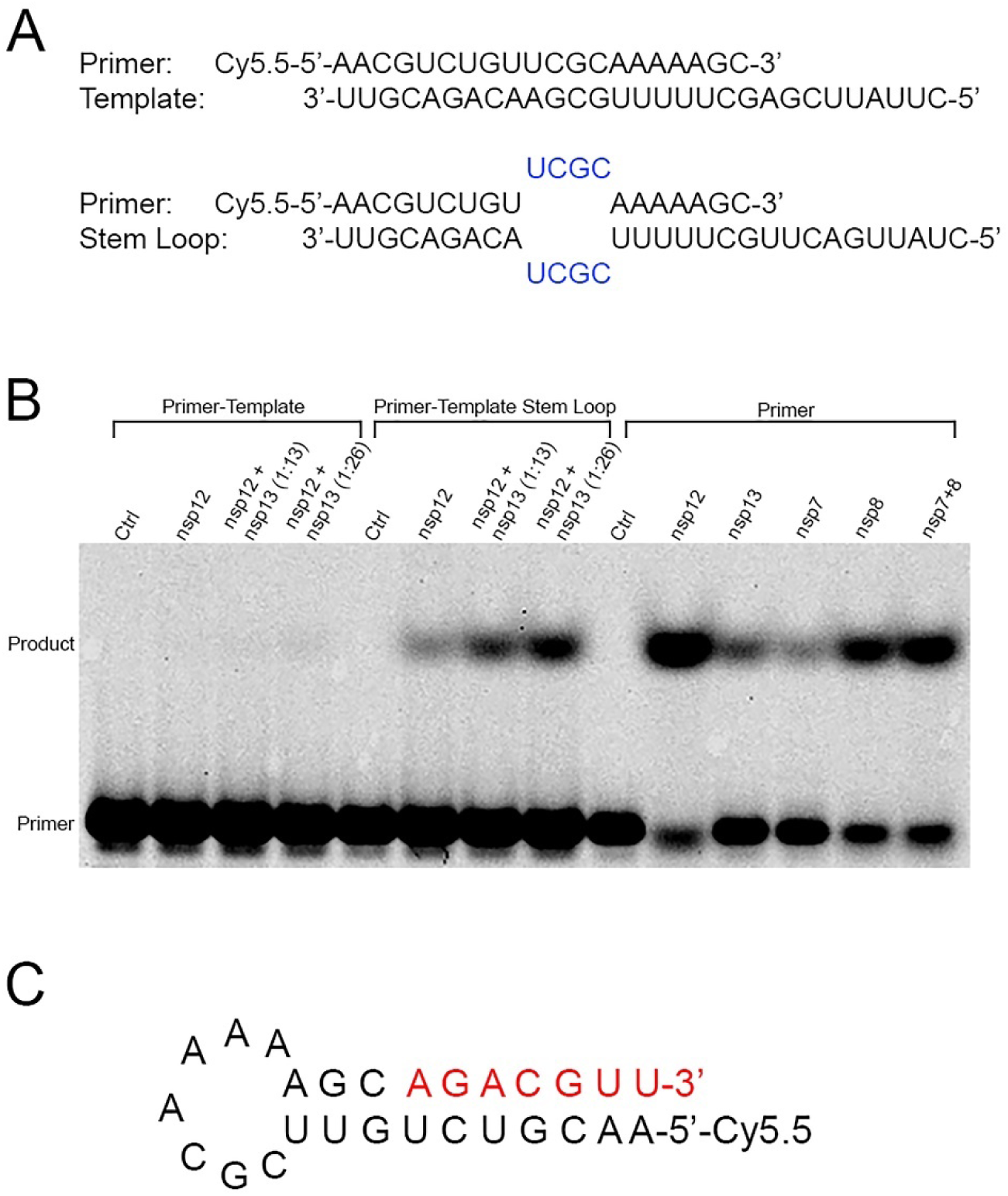
Back-priming extension activity of nsp12 on single-stranded RNA primer. **(A)** Two RNA templates were used in this study. One perfectly paired with the Cy5.5-labeled RNA primer, which had been used in **Figure 1**. The other one formed a stem-loop structure with the primer. The primer was annealed to template in a ratio of 1:3 to form a P/T complex. **(B)** The extension activity of nsp12 on the two primer-template complexes was determined with combination of SARS-CoV helicase nsp13. Two combination ratios were used: 1:13 and 1:26. The assay contained 25 mM Tris HCl pH 8.0, 50 mM KCl, 1 mM DTT, 1 mM MgCl2, 10 nM P/T complex, 16 nM nsp12, 210 or 420 nM nsp13, 100 μM rNTPs (25 μM each), 10% Glycerol, 6 mM NaCl and 0.02% triton X-100. The activities of nsp12, nsp13, nsp7 and nsp8 on the single-stranded primer without a template were also determined. 50% Glycerol was used as controls. Reaction mix was incubated at 37°C for 2h. **(C)** Back-priming mechanism was proposed. The sequence and secondary strucuture of the extended product was predicted. Primer was shown in black and the extension was shown in red.

### Catalytic metals were required for nsp12 activity

To characterize nsp12, we first performed time-course study to analyze the activity of nsp12 at different time-points using the single-stranded RNA primer **(Figure 3A)**. Extended product could be observed as early as 5 min, suggesting that the RNA extension by nsp12 was very fast. The product increased in a time-dependent manner with the decrease of primer. At 30 min, significant amount of product could be observed. At 120 min, almost full activity was observed with little primer left. This study confirmed the activity of nsp12 observed in **Figure 2B** and suggested the extension of the single-stranded RNA primer by nsp12 was very efficient.

**Figure 3.**
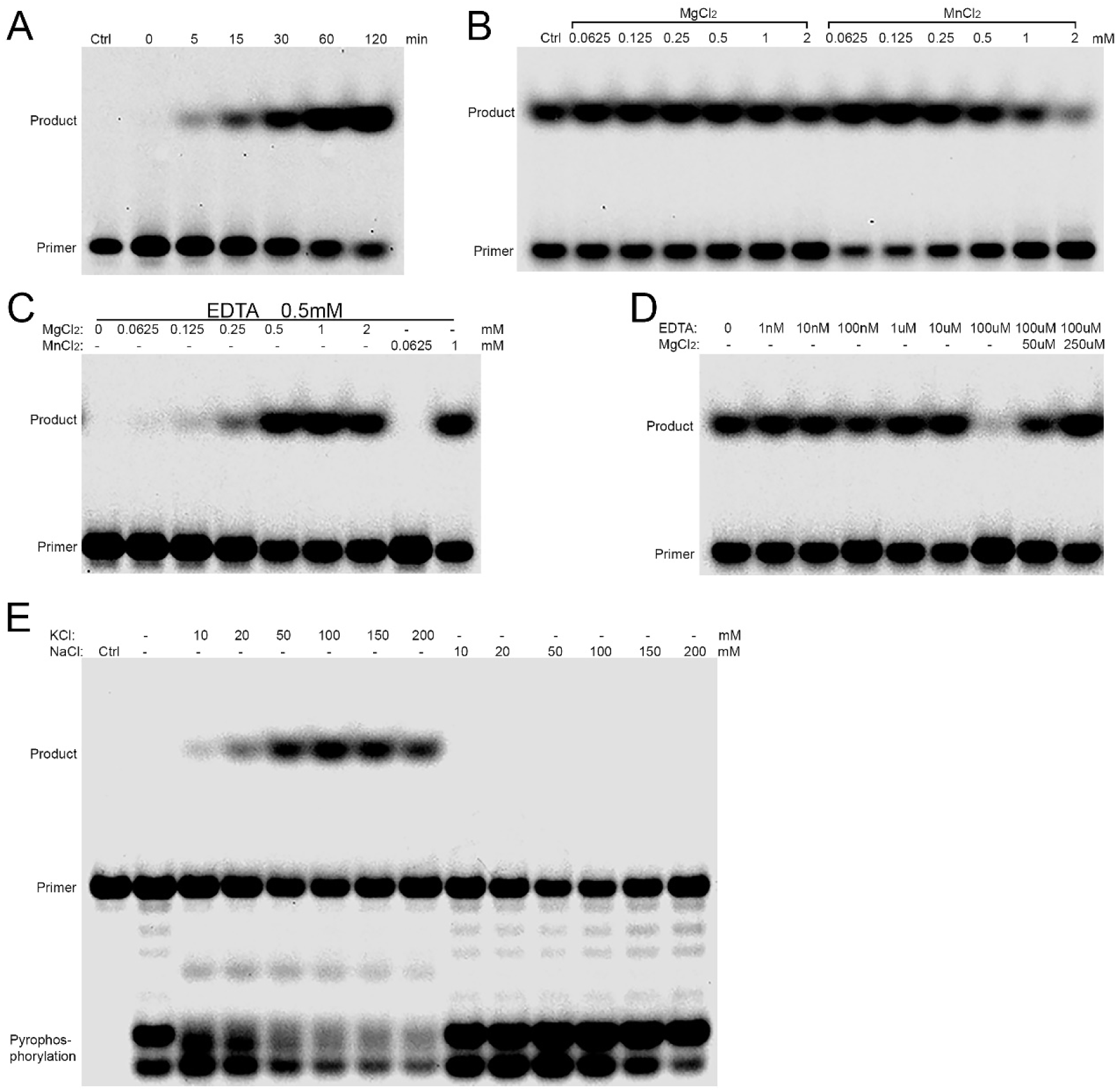
The dependence of nsp12 activity on Mg^2+^ and K^+^. **(A)** Time-course study which showed the activity of nsp12 at various time-points towards the single-stranded RNA primer without a template. The concentration of nsp12 and the RNA primer was 16 nM and 10 nM, respectively. **(B)** showed the activity of nsp12 on the primer under various concentrations of Mg^2+^ or Mn^2+^, from 0.0625 to 2 mM. Control was added with same volume of water. The assay indicated the presence of endogenous catalytic metals, shown by the activity of control. **(C)** The assay was added with 0.5 mM EDTA to block endogenous catalytic metals and the nsp12 activity under various concentrations of Mg^2+^ or Mn^2+^ was re-determined. **(D)** The concentration of endogenous catalytic metals was titrated by serially diluted EDTA. Water was used as control. **(E)** Nsp12 activity was determined at various concentrations of K^+^ or Na^+^, which showed that nsp12 was K^+^-dependent. Pyrophosphorylation activity was observed for Na^+^. 50% Glycerol was used as enzymatic control. The concentration of MgCl_2_ in the assay was 1 mM.

DNA or RNA polymerases required a divalent cation, mostly Mg^2+^, to catalyze DNA or RNA extension. To further characterize nsp12, we determined the dependence of nsp12 on Mg^2+^ and Mn^2+^ **(Figure 3B)**. Both Mg^2+^ and Mn^2+^ could catalyze the extension by nsp12 and optimal activities were observed from 62.5 μM to 1 mM for Mg^2+^ and from 62.5 μM to 0.25 mM for Mn^2+^. Intracellular Mg^2+^ concentration is 0.5-1 mM which was within this range [93] and intracellular Mn^2+^ concentration ranges from 10 to 200 μM [94, 95] which was also widely overlapped with the concentrations evaluated. At high concentrations (2 mM), Mg^2+^ and Mn^2+^ started to decrease the activity of nsp12. This result showed that a wide range of concentrations of Mg^2+^ or Mn^2+^ could support the optimal activity for nsp12. To our surprise, the control without adding Mg^2+^ or Mn^2+^ also showed activity, though sub-optimal. This should be due to the endogenous catalytic metals introduced by protein purification buffers or other reagents used in the assay. To prove this hypothesis, we repeated the assay in the presence of 0.5 mM EDTA **(Figure 3C)**, which could chelate catalytic metals in 1:1 ratio. As expected, activity of the control without Mg^2+^ or Mn^2+^ was blocked completely. The activities of Mg^2+^ at low concentrations (62.5-250 μM) were also blocked while optimal activities were observed from 0.5 to 2 mM. For Mn^2+^, activity was blocked at 62.5 μM and optimal activity was observed at 1 mM. The activity blocking by EDTA suggested that the activity in the control was due to endogenous catalytic metals and nsp12 did require Mg^2+^ or Mn^2+^ for RNA extension. Furthermore, we performed the assay without adding Mg^2+^ or Mn^2+^ and titrated the concentration of endogenous catalytic metals using serially diluted EDTA **(Figure 3D)**. As compared to control without EDTA, activities remained optimal from 1 nM to 10 μM, suggesting EDTA in the range was insufficient to chelate the metals. However, at 100 μM, the activity of nsp12 was fully blocked. As EDTA blocked metals in 1:1 ratio, this result suggested that the concentration of endogenous catalytic metals was below 100 μM. When Mg^2+^ was re-introduced into the assay system, activity was re-observed. Taken together, these results suggested that catalytic metals were required for nsp12 activity and optimal activity could be maintained in a wide range of concentrations of Mg^2+^ or Mn^2+^, including physiological concentrations.

### The activity of nsp12 was K^+^-dependent

Although monovalent cations were not catalytic to DNA/RNA polymerases, they did play an important role in maintaining ion strength which was also essential for polymerase to achieve optimal activities [96, 97]. In addition, monovalent cations also affect the stability of RNA tertiary structure [98-100]. To characterize how monovalent cations would affect nsp12, we determined the activity of nsp12 at various concentrations of K^+^ and Na^+^ **(Figure 3E)**. As compared to the enzymatic control which was performed with 50% glycerol instead of nsp12, RNA extension activities were observed for K^+^, but not for Na^+^. The extension activities increased in a dose-dependent manner from 10 to 50 mM and maintained optimal from 50 to 150 mM. Physiological concentration of K^+^ was 140-150 mM which was within this range [101]. At 200 mM, the activity started to decrease. In contrast, for Na^+^, no extension activity was detected throughout the concentrations evaluated (10 to 200 mM). However, abundant pyrophosphorylation products were observed, indicating that nsp12 was also active in the presence of Na^+^ but the activity only led to pyrophosphorylation. In addition, the control without adding K^+^ or Na^+^ which contained 6 mM NaCl introduced by nsp12 and Cy5.5-RNA stocks, also showed pyrophosphorylation. This result suggested that nsp12 depended on K^+^ to have RNA extension activity and optimal activity required high K^+^ concentration.

### Discovery of 10 nucleotide analogs with inhibition against SARS-CoV nsp12

Based on the characterization of nsp12, we then developed a competitive assay for drug screening. The assay contained 25 mM Tris HCl pH 8.0, 50 mM KCl, 1 mM DTT, 1 mM MgCl_2_, 16 nM nsp12, 10 nM single-stranded RNA primer, 100 μM rNTPs (25μM each) and a nucleotide analog. The analog would compete with its corresponding rNTP, either ATP, UTP, GTP or CTP, and inhibit the RNA extension by nsp12. We collected a small drug library which contained 24 nucleotide analogs, mostly FDA-approved, including two gemcitabine from different suppliers. The analogs were all in their corresponding triphosphate forms to be active in biochemical assays. But to simplify, they were represented by their common drug names instead of triphosphates, unless otherwise specified. Initially, 12 of the analogs were tested as a trial and we found 2’-C-M-GTP could inhibit the RNA extension by nsp12 at 2 mM **(data not shown)**. To optimize the concentration for screening, we then performed dose-response study for 2’-C-M-GTP **(Figure 4A)**. Control was added with 100 μM rNTPs and same amount of water. From 0.25 to 1 mM, 2’-C-M-GTP showed similar levels of product to the control, suggesting it did not inhibit nsp12 at these concentrations. At 2 mM, inhibition could be observed as shown by the decrease of product. And at 4 mM, nearly full inhibition was observed. Inhibition on pyrophosphorylation was also observed for 2’-C-M-GTP at 4 mM. Remdesivir at 2 mM was not effective. This assay showed less products with full extension instead of similar amount of products with shorter extension. This type of inhibition was surprising as nucleotide analogs were supposed to be incorporated into extending RNA and terminate further chain extension, thus generating shorter products. Assays for the evaluation of chain termination were usually non-competitive, in which a nucleotide analog was present but its corresponding natural rNTP, either ATP, UTP, GTP and CTP was absent, which forced the incorporation of the analog. In our competitive assay, all 4 rNTPs were present. Therefore, the generation of fully extended product was possible, even in the presence of a nucleotide analog. These nucleotide analogs might have interfered the RNA extension by nsp12 without incorporation. Or they could be removed from the RNA chain upon incorporation by the pyrophosphorylation activity of nsp12 which had been clearly shown above. Both could slow down the extension process and cause the decrease of product. The competitive assay also resembled better the intracellular condition during drug treatment in which all 4 rNTPs were present as well.

**Figure 4.**
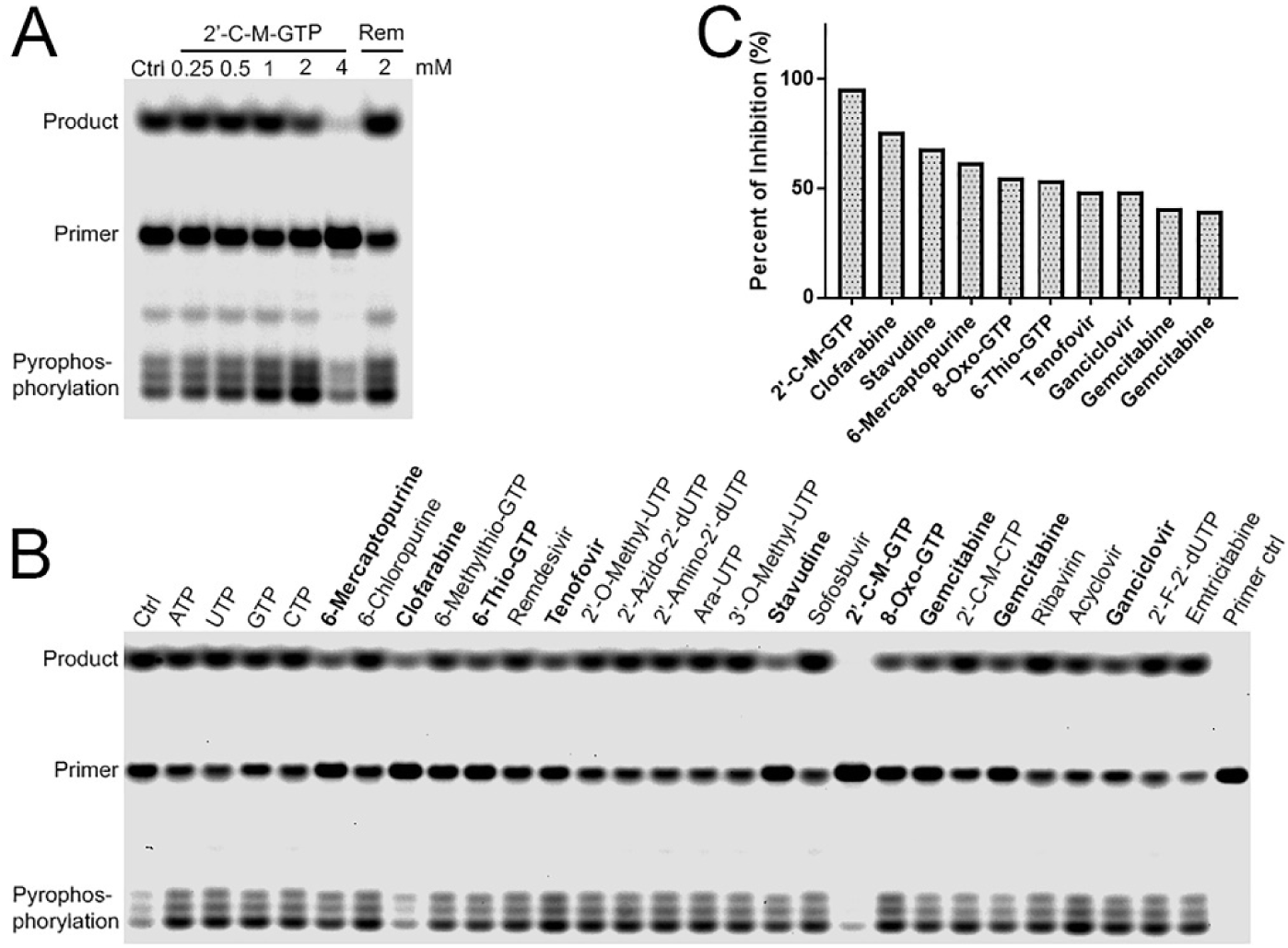
Screening of 24 nucleotide analog triphosphates with inhibition on nsp12. A competitive assay was developed which contained 16 nM nsp12, 10 nM RNA primer, 100 μM rNTPs (25 μM each) and a nucleotide analog in its corresponding triphosphate form. Reaction mix was incubated at 37°C for 30 min. An analog with inhibition on nsp12 would compete with a rNTP and decrease the amount of extended RNA product. **(A)** showed the dose-response of 2’-C-M-GTP and complete inhibition was observed at 4 mM which was used for the following screening concentration. Water was used as negative control. Remdesivir at 2 mM was also evaluated which showed no inhibition. **(B)** The inhibition of 24 nucleotide analogs in their corresponding triphosphate forms were screened at 4 mM using the competitive assay. Emtricitabine was screened at 2 mM due to low stock concentration. Same concentration of primer was loaded as negative control. 100 μM rNTPs without adding extra nucleotide analog was also included as assay control. ATP, UTP, GTP and CTP at 4 mM was used as normalization controls and their activities were averaged and normalized to 100%. Relative percent of inhibition was calculated and hits with >40% inhibition were highlighted in bold. Inhibition on pyrophosphorylation was also observed for clofarabine and 2’-C-M-GTP. **(C)** showed percent of inhibition of the hits.

Based on the dose-response study of 2’-C-M-GTP, 4 mM was chosen as the screening concentration for the 24 nucleotide analogs, except for emtricitabine which was screened at 2 mM due to its low stock concentration. ATP, UTP, GTP and CTP at 4 mM were used as controls. Their activities were averaged and defined as 100%. Relative activity upon nucleotide analog treatment was normalized to the average and percent of inhibition was calculated. **Figure 4B** showed the extended products by nsp12 upon treatment with the 24 nucleotide analogs. **Figure 4C** showed the percent of inhibition of the top 10 hits in the screening. Full list of inhibition by the 24 nucleotide analogs was shown in **Table 1**. Compared to rNTPs control, 10 nucleotide analogs showed >40% inhibition and 14 showed >20% inhibition including remdesivir, as shown by the decrease of extended products. Interestingly, the two gemcitabine from different suppliers showed identical levels of inhibition which suggested the screening assay had high consistency. Of the 10 hits with >40% inhibition, 6 were anticancer drugs, including clofarabine, 6-mercaptopurine, 8-oxo-GTP, 6-thio-GTP and the 2 gemcitabine, and 4 nucleotide analogs were antiviral drugs, including 2’-C-M-GTP, stavudine, tenofovir and ganciclovir. All the 10 nucleotide analogs, except for 2’-C-M-GTP and 8-oxo-GTP, were FDA-approved. And the 3 FDA-approved antiviral hits, stavudine, tenofovir and ganciclovir, had acceptable safety profiles. Emtricitabine, ribavirin and sofosbuvir, which are currently under evaluation for COVID-19 in clinical trials, showed minimal inhibition against SARS-CoV nsp12. They either did not inhibit nsp12 or required higher concentration to be effective. Or, they could have involved other mechanisms to inhibit virus replication. This has been reported for ribavirin which can inhibit virus replication by introducing mutations instead of inhibiting RNA/DNA replication [102]. The effective concentration of ribavirin in cell culture could also be high for some viruses [82, 103]. In addition, as compared to rNTP controls, the pyrophosphorylation products of clofarabine and 2’-C-M-GTP were decreased while most of the nucleotide analogs showed similar levels. As the top 2 hits, their higher inhibition might have led to the decrease of pyrophosphorylation, or they might indicate a different mechanism to inhibit RNA extension.

**Table 1.** Inhibition of 24 nucleotide analogs on SARS-CoV nsp12 and nsp8.

| Order | Nucleotide<br>Analogues | Inhibition on<br>Nsp12 (%) | Inhibition on<br>Nsp8 (%) | Specificity to<br>Nsp12 (%) |
| --- | --- | --- | --- | --- |
| 1 | Primer control | 100.00 | 100.00 | 0.00 |
| 2 | 2'-C-M-GTP | 95.94 | 98.63 | -2.69 |
| 3 | Clofarabine-TP | 75.84 | 62.27 | 13.57 |
| 4 | Stavudine-TP | 68.16 | 44.60 | 23.56 |
| 5 | 6-Mercaptopurine-TP | 62.05 | 59.94 | 2.11 |
| 6 | 8-Oxo-GTP | 55.38 | 69.45 | -14.07 |
| 7 | 6-Thio-GTP | 53.77 | 53.62 | 0.15 |
| 8 | Tenofovir-DP | 48.94 | 47.24 | 1.71 |
| 9 | Ganciclovir-TP | 48.94 | 33.74 | 15.20 |
| 10 | Gemcitabine-TP | 40.90 | 73.56 | -32.65 |
| 11 | Gemcitabine-TP | 40.10 | 70.43 | -30.33 |
| 12 | 6-Methylthio-GTP | 30.85 | 23.31 | 7.54 |
| 13 | Remdesivir-DP | 28.44 | 1.23 | 27.22 |
| 14 | Acyclovir-TP | 26.03 | 35.58 | -9.55 |
| 15 | 2'-O-methyl-UTP | 21.21 | 9.20 | 12.00 |
| 16 | 2'-Amino-2'-dUTP | 14.37 | -9.20 | 23.57 |
| 17 | 2'-Azido-2'-dUTP | 10.35 | -12.88 | 23.24 |
| 18 | Emtricitabine-TP | 10.35 | 15.34 | -4.99 |
| 19 | ATP | 7.54 | -4.91 | 12.45 |
| 20 | 2'-C-M-CTP | 5.53 | 22.09 | -16.56 |
| 21 | 2'-F-2'-dUTP | 5.13 | -2.45 | 7.58 |
| 22 | Ara-UTP | 4.72 | 21.47 | -16.75 |
| 23 | CTP | 1.51 | -4.29 | 5.80 |
| 24 | 3'-O-methyl-UTP | -0.10 | -1.23 | 1.13 |
| 25 | GTP | -0.90 | 13.50 | -14.40 |
| 26 | 6-Chloropurine-TP | -2.91 | -15.34 | 12.42 |
| 27 | 100µM rNTP ctrl | -7.34 | -15.34 | 8.00 |
| 28 | UTP | -8.14 | -5.52 | -2.62 |
| 29 | Ribavirin-TP | -24.62 | -12.88 | -11.74 |
| 30 | Sofosbuvir-DP | -27.44 | -21.47 | -5.96 |

The top 10 hits were verified at 1, 2 and 4 mM, using the same assay as screening **(Figure 5A)**. All hits showed significant decrease of extension product in a dose-dependent manner, except for ganciclovir which showed similar level of extension product to ATP control at 4 mM. In addition, as an approved treatment for COVID-19, remdesivir was also included in the verification assay which confirmed its inhibition on nsp12 at 4 mM as observed in the screening. Inhibition on pyrophosphorylation for clofarabine and 2’-C-M-GTP at 4 mM was also confirmed. Taken together, this study supported the validity of the screening and confirmed all the hits in the screening, except for ganciclovir, had obvious inhibition on nsp12.

**Figure 5.**
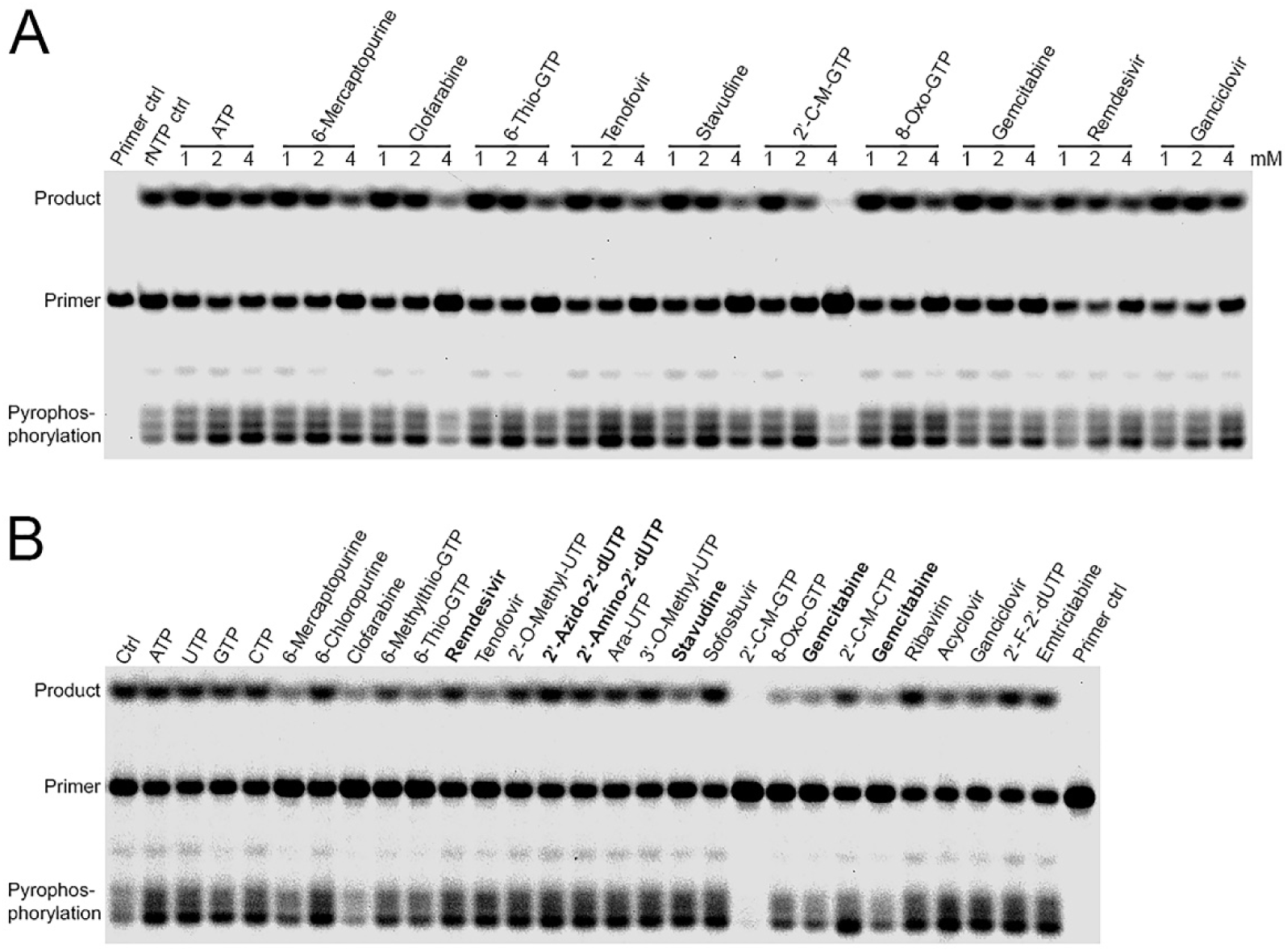
Verification of nsp12 hits and the screening of nsp8. **(A)** The nsp12 hits with >40% inhibition were verified at 1, 2, and 4 mM using the same assay as screening. Dose-dependent decrease of extended RNA product was observed. Remdesivir which had 28.44% inhibition on nsp12 was included as well. ATP was used as normalization control. Same concentration of primer was used as primer control. 100 μM rNTPs without adding extra nucleotide was used as rNTP control. **(B)** The inhibition on nsp8 of the 24 nucleotide analog triphosphates were also screened to identify nsp12-specific inhibitors and their inhibition percentages on nsp8 were compared to nsp12. Drugs with >20% differential inhibition between nsp12 and nsp8 were highlighted in bold.

### Discovery of stavudine and remdesivir as nsp12-specific antivirals

As the competitive screening assay developed in this study was novel, we tried to prove the validity of the assay from different perspectives. One good way is probably to prove the hits in the screening were specific to nsp12. In **Figure 2B**, nsp8 was shown to possess comparable RNA extension activity to nsp12, although it was tested at a much higher concentration. Nsp8 had been proposed as a primase and certain mutations in nsp8 region could lead to replication deficiency, even lethality, for SARS-CoV [88]. Nsp8 was 3-fold smaller in size than nsp12 and lacked the seven conservative motifs A to G, making nsp8 a reasonable control to identify nsp12-specific drugs. We performed screening against nsp8 using the 24 nucleotide analogs **(Figure 5B)**. The hits and their specificity to nsp12 were listed in **Table 1**. To our surprise, most of the analogs in nsp8 screening showed similar patterns of inhibition to nsp12. This result either suggested that the inhibition of RNA extension by these nucleotide analogs was independent of nsp12 or nsp8, or suggested that nsp12 and nsp8 might have evolved similar composition in the catalysis active site to better coordinate the RNA replication for SARS-CoV. Interestingly, we did observe that the inhibition of stavudine decreased by 20% from nsp12 to nsp8 and the inhibition of gemcitabine increased by 30%, suggesting that stavudine had more specificity towards nsp12 while gemcitabine had less. In addition, remdesivir, which had 28.44% inhibition against nsp12, showed minimal inhibition against nsp8, suggesting that remdesivir was a nsp12-specific drug completely. The specificity of stavudine and remdesivir to nsp12 did support the hits validity of the screening. To note, nsp12 is the major RdRp for SARS-CoV and a specific drug may be favorable as they could improve efficacy and reduce side effects. However, the efficacy can also be enhanced if a drug had dual inhibition on nsp12 and nsp8. Therefore, the comparison between nsp12 and nsp8 screenings was only to prove the validity of the screening and not necessarily to be able to identify a more potent hit.

### Stavudine, tenofovir, clofarabine and gemcitabine had inhibition on 2019-nCoV nsp12

Nsp12 was highly conservative among SARS-CoVs, inhibitors of SARS-CoV nsp12 most likely would inhibit 2019-nCoV as well. To confirm, we purified the nsp12 of 2019-nCoV and evaluated its activity by the same single-stranded RNA primer used for SARS-CoV. As shown by time-course study **(Figure 6A)**, extended product could be detected as early as 5 min and full activity was achieved at 120 min. This result suggested that the nsp12 of 2019-nCoV could extend the RNA primer as efficiently as SARS-CoV. Based on potency and safety, we then selected 4 of the 10 SARS-CoV hits, stavudine, tenofovir, clofarabine and gemcitabine, in their corresponding triphosphates, to be evaluated on 2019-nCoV nsp12 **(Figure 6B-C)**. Ganciclovir and remdesivir were also included. As compared to ATP control, stavudine, tenofovir, clofarabine and gemcitabine showed significant decrease of extension products, while ganciclovir and remdesivir showed low levels of decrease. This result suggested that stavudine, tenofovir, clofarabine and gemcitabine could inhibit 2019-nCoV nsp12 as well.

**Figure 6.**
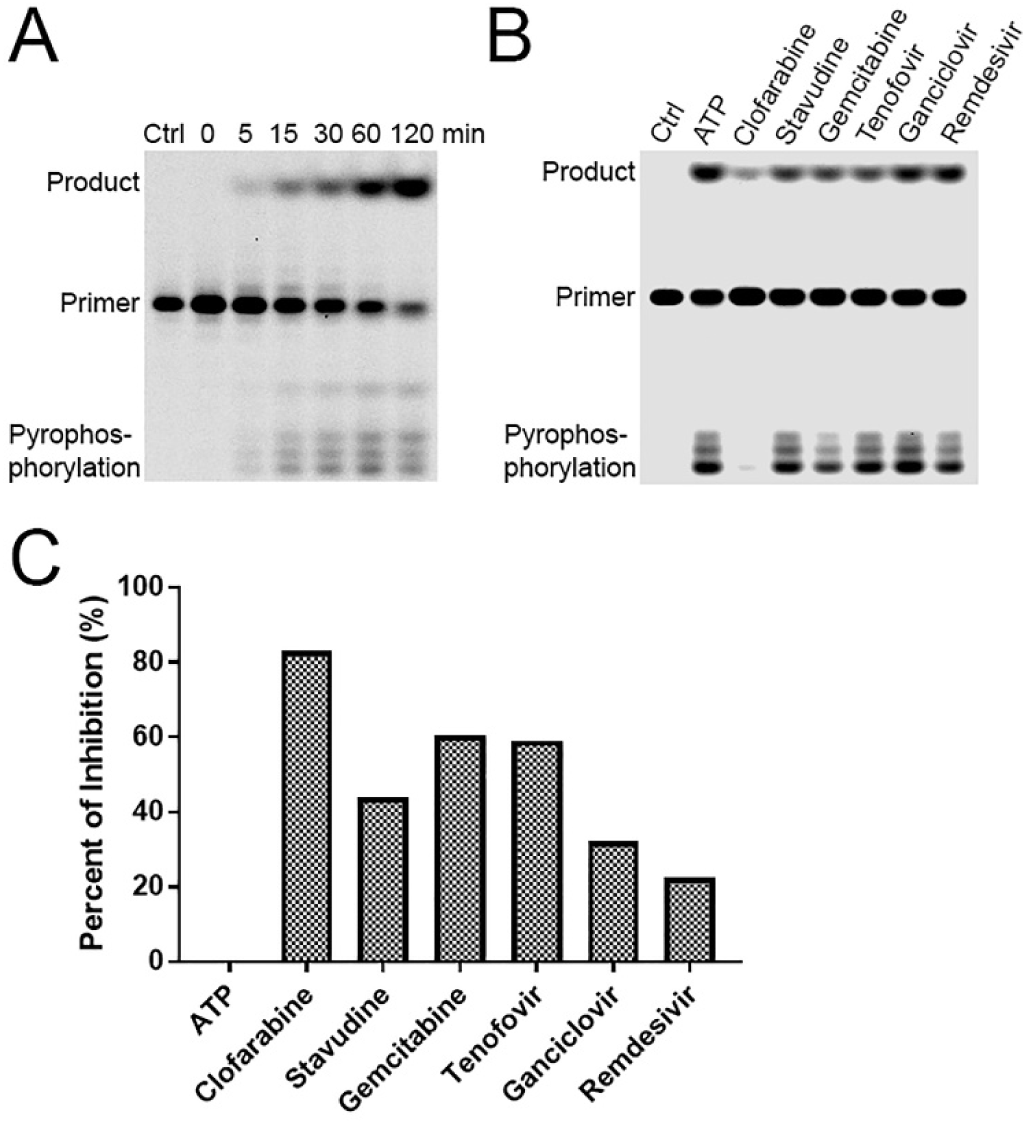
The inhibition of selected SARS-CoV hits on 2019-nCoV nsp12. **(A)** The activity of 2019-nCoV nsp12 was determined at different time-points which showed similar activities to SARS-CoV. 50% Glycerol incubated for 120 min was used as control. **(B)** Clofarabine, stavudine, gemcitabine, tenofovir, ganciclovir and remdesivir from SARS-CoV screening were selected and their inhibition on 2019-nCoV nsp12 was evaluated at 4 mM which showed inhibition as well. The assay condition was same as SARS-CoV. The analogs were evaluated in their corresponding triphosphate forms. ATP at 4 mM was used as normalization control. Same concentration of primer was loaded as negative control. **(C)** showed the percent of inhibition as compared to ATP of which the activity was defined as 100% and inhibition as zero.

### Discovery of five HIV nucleoside analog reverse-transcriptase inhibitors (NRTIs) with inhibition on nsp12

HIV NRTIs had good safety profiles, making them ideal candidates for drug re-purposing. In the screening, 3 HIV NRTIs were evaluated and 2 of them, stavudine and tenofovir, showed inhibition on nsp12. It seemed HIV NRTIs had high possibility to inhibit the nsp12 of SARS-CoVs, though the sample size was very small. To date, 8 NRTIs in total have been approved for HIV treatment. Five of them were not included in the screening library due to commercial availability or delivery issues. We managed to purchase all 8 NRTIs and tested them on nsp12 in their corresponding active triphosphates. Clofarabine, an anticancer drug, was included as treatment control. A primer control without nsp12 was also included. As shown by **Figure 7A**, in addition to stavudine (44.64%) and tenofovir (43.23%), another 3 NRTIs, abacavir, zidovudine and zalcitabine, were identified to be effective inhibitors against SARS-CoV nsp12, with 50.09%, 34.62%, and 89.67% inhibition, respectively **(Figure 7C)**. The nsp12 of 2019-nCoV showed identical results **(Figure 7B)**, with 53.05%, 37.56% and 84.69% inhibition for the 3 NRTIs, respectively **(Figure 7C)**. The inhibition by zalcitabine was almost complete at 4 mM. In addition, though not regarded as a hit, lamivudine also showed low level of inhibition on both viruses (20.39% and 15.02%, respectively). To verify the inhibition of the 3 newly identified NRTIs, they were evaluated at three concentrations (1, 2 and 4 mM), with ATP and tenofovir used as controls **(Figure 8)**. The inhibition of all 3 NRTIs were confirmed at 4 mM and zalcitabine also showed inhibition at 2 mM. Taken together, these results suggested that 5 HIV NRTIs, tenofovir, stavudine, abacavir, zidovudine and zalcitabine, were effective inhibitors against the nsp12 of both SARS-CoV and 2019-nCoV.

**Figure 7.**
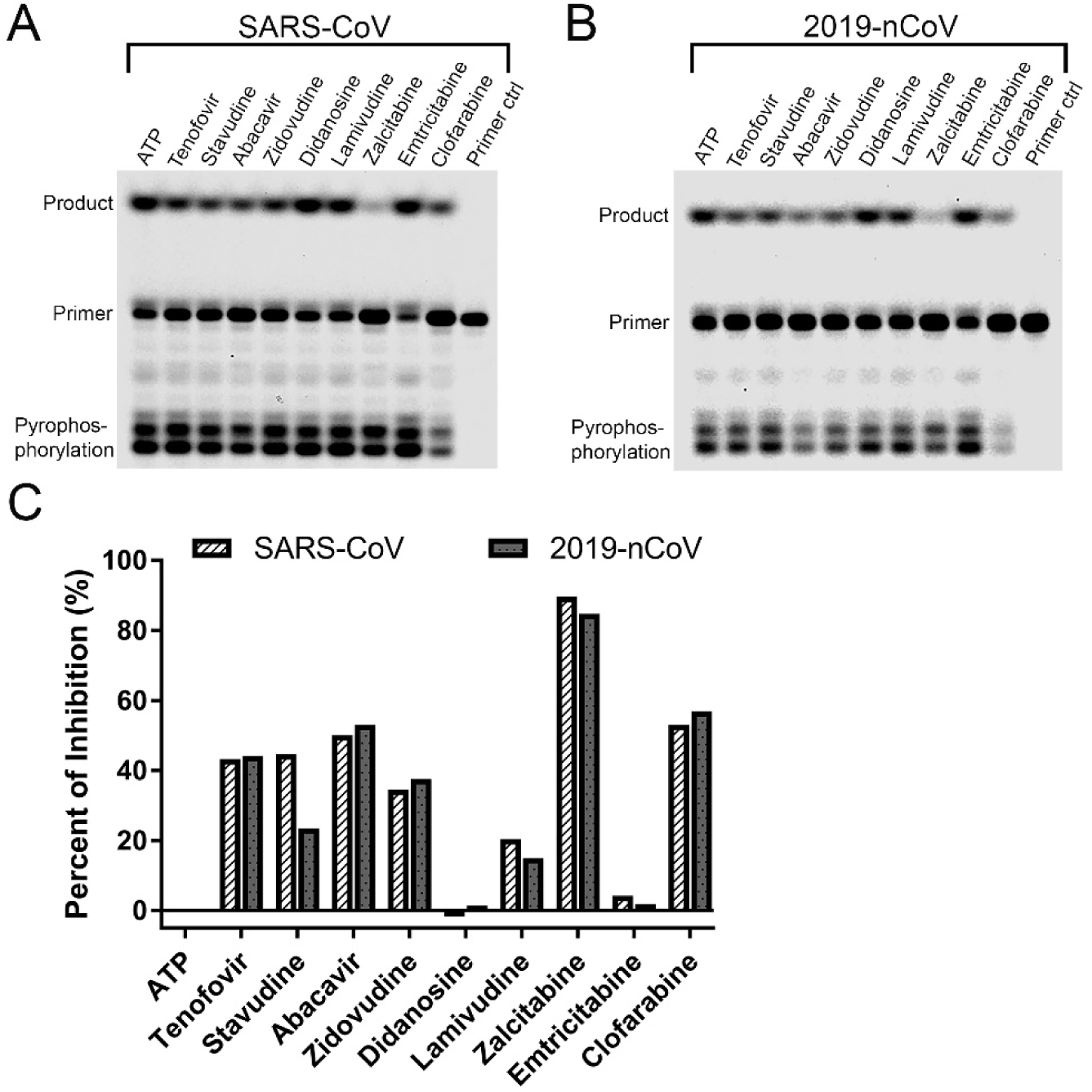
Drug-repurposing of HIV nucleoside analog reverse-transcriptase inhibitors (NRTIs) as antivirals against SARS-CoV and 2019-nCoV. Eight FDA-approved HIV NRTIs were evaluated for their inhibition on the nsp12 of SARS-CoV **(A)** and 2019-nCoV **(B)**. The assay contained 16 nM nsp12, 10 nM RNA primer, 100 μM rNTPs and 4 mM a NRTI, in its corresponding active triphosphate form. The extended product of the RNA primer by nsp12 upon NRTI treatment was analyzed by Urea-PAGE. Product was quantified and percent of inhibition was calculated. ATP was used as normalization control. Clofarabine, an anticancer drug, was used as treatment control. Assay with 50% glycerol instead of nsp12 was used as primer control. **(C)** showed the percent of inhibition of the 8 NRTIs, compared to ATP control. SARS-CoV was presented by white bar with slashes. 2019-nCoV was presented by gray bar with dots.

**Figure 8.**
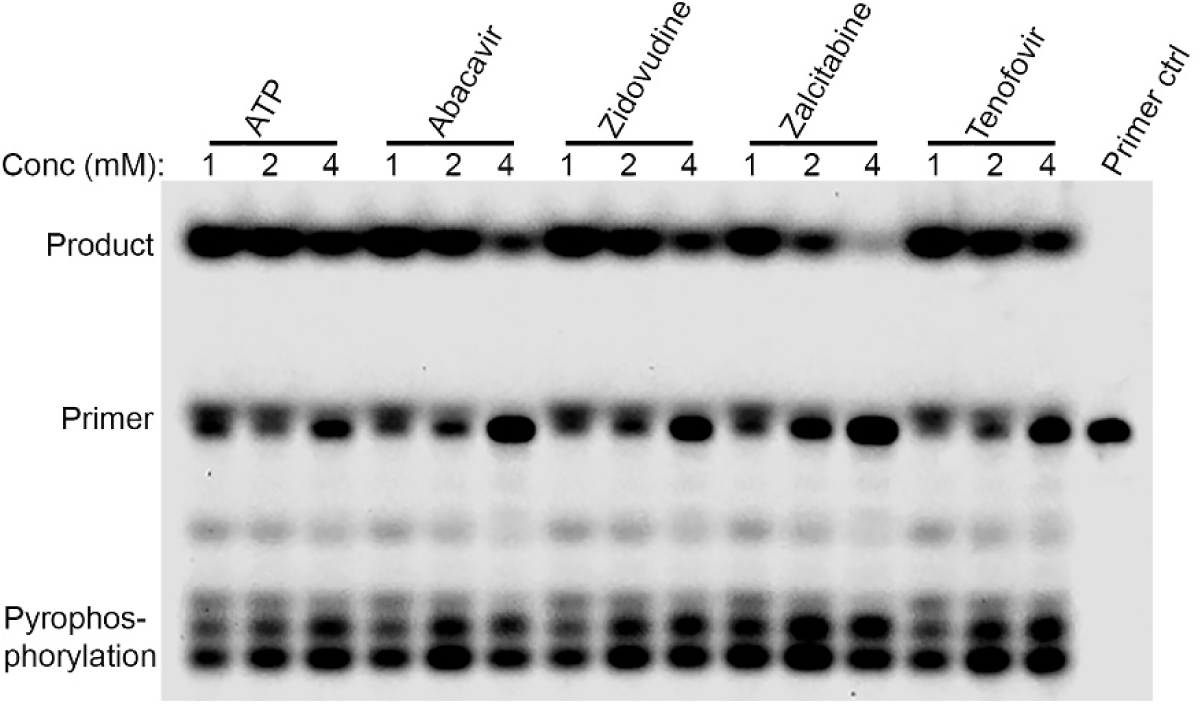
Inhibition verification of abacavir, zidovudine and zalcitabine on nsp12. The three newly identified NRTIs, abacavir (carbovir-TP), zidovudine and zalcitabine triphosphates, were verified at three concentrations (1, 2 and 4 mM) for their inhibition on the RNA extension by nsp12, using SARS-CoV as a model. The Cy5.5-labeled RNA primer was used and its extended product as well as pyrophosphorylation product by SARS-CoV nsp12 were visualized. ATP was used as treatment negative control. Tenofovir was used as treatment positive control. Reaction with 50% glycerol instead of nsp12 was used as primer control. For abacavir and zidovudine, inhibition was confirmed at 4 mM. For zalcitabine, inhibition was observed at both 2 and 4 mM.

## Discussion

### The dependence of nsp12 activity on K^+^ and divalent catalytic metals

In this study, we expressed and purified active SARS-CoV nsp12 which could efficiently extend a single-stranded RNA. The activity depended on K^+^ while Na^+^ led to pyrophosphorylation and optimal activity was observed at concentrations close to the physiological K^+^ concentration (140 to 150 mM) [101]. To our knowledge, this is the first time such dependence had been reported for nsp12. Previously, nsp12 alone was reported to be inactive [65, 88], which possibly could be explained by the conditions in which the activities were determined, either with Na^+^ or low concentration of K^+^. In our study, the activity of nsp12 was also dependent on Mg^2+^ or Mn^2+^, but due to its intracellular predominance [93], Mg^2+^ should be the major catalytic metal for nsp12 physiologically. Interestingly, Mn^2+^ is mutagenetic and promotes mis-incorporation during RNA synthesis. Viral RdRps in general had low replication fidelity and a relative high mutation rate. Therefore, it is possible that nsp12 may take advantage of Mn^2+^ as a cofactor to enhance mutation during replication by occasionally involving it into the catalysis active site. Furthermore, our study proved that nsp12 alone could be fully active at low concentration (16-32 nM) without the requirement of nsp7 and nsp8. The full activity was only observed for single-stranded RNA, possibly via a back-priming mechanism [88]. While for double-stranded RNA, the activity of nsp12 was very low which was consistent with previous studies [58, 88]. As single-stranded RNA was unstable, it may have intrinsic drive to back-prime and extend to form a stable double-stranded structure. This intrinsic drive would promote the extension by nsp12 which could explain the low activity of nsp12 observed for double-stranded RNA.

### Competitive assay for antiviral discovery against nsp12

To discover nucleotide analogs against nsp12, a competitive assay was developed which provided a rapid screening method to identify drugs with inhibition on nsp12. Assays in previous studies were usually based on chain termination in which the corresponding rNTP of a nucleotide analog, either ATP, UTP, CTP or GTP, was absent to determine the efficiency of the analog to be incorporated into RNA chain and the efficiency to cause chain termination. In contrast, we developed a competitive assay. All the 4 rNTPs were present as well as a nucleotide analog. The nucleotide analog would compete with its corresponding rNTP, which resembled the intracellular condition during nucleotide analog treatment. Instead of showing chain termination with shorter product, the assay showed decrease of the fully extended product. As a catalyst, nsp12 could catalyze both RNA polymerization and the reverse process, pyrophosphorylation. Therefore, it was highly possible that nsp12 could remove chain-terminating nucleotide analogs from RNA chain upon incorporation, leading to the absence of a terminated product. To our knowledge, this is the first time that nsp12 has been shown to be associated with such property which provides a new perspective about how nsp12 replicates SARS-CoVs genome intracellularly.

### High concentration for nsp12 antiviral screening

In this study, we performed drug screening and identified 10 nucleotide analogs with >40% and 14 with >20% inhibition on nsp12, including remdesivir, gemcitabine and tenofovir. The screening concentration was 4 mM which was high. This should be due to the short extension (7 bp) of the RNA product and high concentration of nucleotide analogs had to be involved to show inhibition. SARS-CoVs have a genome of 30 kbp and nucleotide analogs would be able to achieve similar levels of inhibition at much lower concentrations intracellularly. Thus, further evaluation in cell culture is necessary to better determine the effective concentrations of these analogs. But for primary screening, this assay is sufficient for hits identification. In the screening, remdesivir showed 28.44% inhibition on nsp12 and its effects has been proven in cell cultures with EC_50_= 0.01 μM [104]. Another hit in our assay, gemcitabine, was also reported to be able to inhibit SARS-CoV, MERS and 2019-nCoV at micromolar range in cell culture [78, 79]. And recently, one study reported that tenofovir, in its tenofovir disoproxil fumarate (TDF) form, inhibited 2019-nCoV at micromolar range in cell culture [80]. In addition, a cohort study of HIV-positive persons receiving antiretroviral therapy revealed that receiving TDF/FTC could lower the risk of 2019-nCoV hospitalization [105]. These studies provide evidence that hits in our screening can inhibit SARS-CoVs at micromolar range in cell culture.

### Drug-repurposing and combination of HIV NRTIs for COVID-19 treatment

In this study, we identified 5 HIV NRTIs to be effective inhibitors against SARS-CoV and 2019-nCoV. Among them, tenofovir, abacavir and zidovudine have superior safety profiles than stavudine and zalcitabine [106, 107]. Zalcitabine has been discontinued since 2006. Abacavir and stavudine share the same sugar backbone with a carbon-carbon double bond. They probably can be used as an alternative to each other. But as abacavir has a superior safety profile, it would be the first choice. Therefore, tenofovir, abacavir and zidovudine are the top 3 hits we would like to propose as the candidates to be further investigated for COVID-19 treatment. In addition, as proven by antiretroviral therapies, drug combination is a powerful approach to improve therapy efficacy and solve drug resistance issues. Therefore, we would also like to propose the combinations of tenofovir, abacavir and zidovudine to be evaluated for COVID-19 treatment. During SARS outbreak, receiving highly active antiretroviral therapy (HAART) was reported to be a protection factor against the virus [108]. In a recent cohort study, taking tenofovir disoproxil fumarate (TDF) combined with emtricitabine was associated with lower risk of 2019-nCoV hospitalization [105]. As emtricitabine showed minimal inhibition on nsp12 in our study, the combination of tenofovir with abacavir and/or zidovudine probably would have better efficacy against COVID-19, if there is no antagonism among them. Lamivudine can also be included into the combinations as it showed about 20% inhibition on nsp12 in our study. Currently, FDA-approved HIV medicines combined by the 4 NRTIs include Epzicom (abacavir/lamivudine), Trizivir (abacavir/lamivudine/zidovudine), Cimduo (lamivudine/tenofovir disoproxil fumarate), and Combivir (lamivudine/zidovudine) [109]. Another choice for combination is type I interferons which can inhibit the replication of 2019-nCoV in cell culture [110] and have showed positive outcomes in clinical trials [56, 111]. To conclude, we proposed 5 HIV NRTIs and their related combinations as the candidates to be investigated for COVID-19 treatment. And we call for open collaboration to get them further evaluated in cell cultures.

## Conclusion

In this study, we expressed and purified active SARS-CoV nsp12 which could efficiently extend single-stranded RNA in a K^+^ and Mg^2+^-dependent manner. We developed a competitive assay for antiviral screening of nucleotide analogs against nsp12 and identified 10 hits with more than 40% inhibition. We also discovered that stavudine and remdesivir were specific antiviral to nsp12. In addition, 5 FDA-approved HIV NRTIs, tenofovir, stavudine, abacavir, zidovudine and zalcitabine, were identified to be effective inhibitors for the nsp12 of both SARS-CoV and 2019-nCoV. And we proposed the 5 NRTIs and their related combinations to be further investigated as the candidates for COVID-19 treatment.

## Acknowledgement

This study was supported by the Bill & Melinda Gates Foundation, Beijing Municipal Government and Tsinghua University.

---

**Figure S1.**
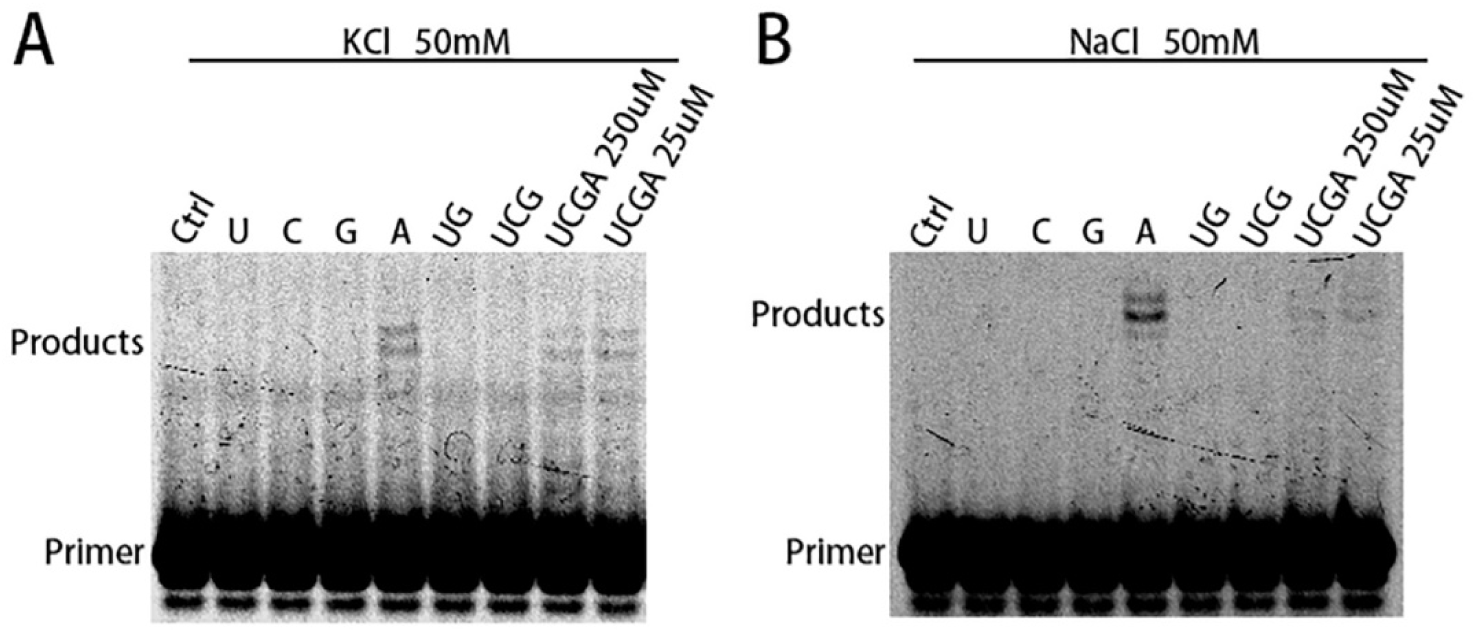
Poly A activity of nsp12 on double-stranded RNA. An RNA primer annealed to a template was used to determine the template-dependent poly A activity of nsp12, with either single rNTP (UTP, CTP, GTP, ATP) or their combinations (UG, UCG, UCGA). The concentration for each rNTP was 250 μM. Water was used as control and a control with 25 μM each of UCGA was also included. The assay was performed with 50 mM either KCl **(A)** or NaCl **(B)**. ATP alone showed identical extended products as UCGA while UTP, CTP, and GTP showed no extension, and the extension length (9 nt) was close to the template. This study suggested nsp12 had poly A activity.

